# Explainable prediction and simulation of complex system dynamics through networks of manifolds

**DOI:** 10.64898/2026.05.12.724527

**Authors:** Joseph Park, Cameron Smith, Shih-Yi Tseng, Jennifer Guidera, Andrei V Semenov, Stanislav Smirnov, Loren M Frank, Gerald M Pao

## Abstract

Complex systems such as brains and other interacting biological and physical processes are difficult to represent because they evolve across many variables, scales, and nonlinear interactions. To capture these multivariate, multiscale interactions we have developed Generative Manifold Networks (GMNs) a machine learning framework consisting of a network of linked dynamical systems. The network is discovered by an interaction function which can focus on causality, shared information, nonlinearity or other metric. Network nodes are low–dimensional data–driven state–space manifolds with generator functions accommodating multiscale dynamics. In contrast to many machine learning approaches GMNs have no latent or randomly initialized variables providing transparent explainability. GMNs generate short term dynamics of chaos on par with echo state networks while outperforming them in long term generation of chaos and neural dynamics, but with a markedly reduced number of dimensions and without sensitive dependence on reservoir parameters or random states. As a result of their holistic, multiscale representation GMNs can learn the complete dynamics of a complex system. We further show that GMNs are universal approximators. GMNs are demonstrated on chaotic dynamics, neural and behavioral recordings of the fruit fly and domestic rat with comparisons to echo state networks and crossformer – a time series transformer.

**Significance:** A major challenge in machine learning is to model complex systems accurately without losing interpretability. Many methods that succeed in prediction rely on latent variables obscuring mechanistic insight and complicating experimental testing. Generative manifold networks (GMN) construct a network of low-dimensional functional manifolds directly from observed variables with no latent or randomly initialized variables: the model remains transparent and experimentally testable. We prove that GMN are universal approximators showing that high representational power can be achieved without sacrificing explainability. GMN therefore provides a general framework for prediction and simulation in neuroscience and complex systems where unraveling the links between variables in an experimentally testable manner is as important as forecasting their behavior.

## 1 Introduction

A central problem in modern data science is how to model complex dynamical systems in a form that is simultaneously expressive, generative, and interpretable. This problem is especially acute in neuroscience, where large–scale recordings reveal highly complex activity, yet neural population dynamics and behavior often evolve on much lower–dimensional structure [1] reflecting the *manifold hypothesis* [2] observed across living systems [3] and especially in neural processing [4, 5]. At the same time, neural function is fundamentally networked arising from interactions across cells, regions, and timescales [6–8]. Similar combinations of low–dimensional structure and network organization are observed across many complex systems, including genomic [9, 10], metabolic [11, 12], physiologic [13], and social systems [14].

It is therefore not surprising that manifolds and networks have become central organizing ideas in machine learning. Indeed, one may view much of the recent success of machine learning as arising from their synthesis, consistent with the unreasonable effectiveness of deep learning [15]. Deep learning has been unusually successful in domains governed by highly nonlinear relationships, including problems for which traditional analytical or physics–based approaches have struggled to produce accurate predictive solutions. In that sense, these successes suggest that useful solutions exist even when they are difficult to derive in closed mechanistic form. However, most such models achieve expressive power by transforming observations into latent variables, hidden states, or through massively overparameterized representations. Although effective for prediction, these transformations often weaken the connection between model components and measured observables, complicating mechanistic interpretation, causal inference, and experimental validation.

For example, low–dimensional manifolds expressed in terms of global latent objects such as principal components or learned latent embeddings can summarize neural population activity, but typically do not preserve an explicit mapping from local observed variables—such as neurons, voxels, or regions of interest (ROI)—to the dynamical structure of the model [16, 17], nor do they naturally encode the network interactions through which local components influence one another across multiple timescales. As successful as low–dimensional manifold representations have been they often remain abstract, not directly observable descriptions that complicate experimental manipulation.

To address these issues, we depart from the prevailing view of neural and behavioral manifolds as single latent objects and instead represent complex dynamics as a network of local low–dimensional manifolds anchored directly to observables. This architecture is termed *generative manifold networks* (GMN). The network is discovered through an interaction function between observables defining an adjacency matrix from which a network graph is grown from a selected observable. The interaction function can be chosen to emphasize the property of interest, including causality through convergent cross mapping (CCM) [18], shared information through mutual information, nonlinear dependence [19–21], or other suitable interaction metrics. Each node of the network is a multivariate state–space manifold constructed through generalized Takens embedding of the relevant variables [22]. Importantly, each node/manifold corresponds directly to a system observable and its interacting variables. The architecture is therefore low–dimensional, data–driven, and observable, enabling prediction and generation of dynamics at all nodes of the network.

These features address several outstanding issues in machine learning and scientific modeling. First, GMN address the growing need for explainability [23] as many statistical and machine learning algorithms [24] transform observations into latent, unobservable spaces that complicate direct hypothesis testing and experimental manipulation. Second, GMN allows network construction to reflect the dynamical relation of interest, helping distinguish correlative from potentially causal links among variables [25]. Third, GMN generates all observable components of a multivariate networked system rather than predicting isolated outputs [26]. Most importantly, we prove that GMN are universal approximators showing that direct observability and explainability need not come at the cost of representational power.

Although GMN was conceived to model neural systems, its contribution is algorithmic and general. We demonstrate GMN on chaotic dynamics, *Drosophila* calcium imaging with behavior, and *Rattus norvegicus* electrophysiology with behavior with results compared to echo state networks (ESN) and a state–of–the–art multivariate time series transformer, Crossformer. Across these settings, GMN provides accurate prediction and simulation of complex dynamics while preserving a direct link between model structure and experimentally accessible observables.

### 1.1 GMN architecture

The GMN architecture is simple: a network of nodes, each node a dynamical system generating time series of an observable from a multivariate generalized embedding of observations. Here, we use the state space simplex generator defined by Sugihara [27] avoiding statistical model constraints such as independence and linearity, however, other generators can be used. The generalized embedding contains multivariate inputs from other nodes while allowing time delayed versions of observed variables to represent unobserved dynamics via Takens theorem [28]. Time delay components are determined by node-specific embedding dimension *E* and time delay *τ* . However a node may have time delay embedding dimension *E* = 1 resulting in no delays, or may have no external inputs incorporating only past observations of the node output.

Importantly, manifolds are entirely data–driven and observable as any delayed components representing unobserved dynamics maintain direct mapping to observations. The network is inherently multiscale according to node embedding parameters and network structure. Network topology can be tailored to examine a specific dynamical aspect of interest through choice of the intervariable interaction function, for example, causality, shared information, synchronized dynamics, or, the network can be manually constructed and manipulated to examine the influence of specific interactions (supplement D).

As an example consider the Lorenz’63 system with three observables V1,V2,V3 shown in figure 1. We designate observable V3 as the network terminus (target). Network discovery proceeds by ranking interaction matrix row V3 (figure 1c bottom row). Starting at the highest ranked node, nodes driving M_V3_ are added as long as no network cycles are introduced, here, nodes M_V2_ and M_V1_ are added as drivers of node M_V3_. Nodes that were added are then recursively explored in the same fashion, the row for V2 is explored adding M_V1_ as a driver of M_V2_. When row V1 is explored there are no new links to be added. The corresponding 3 node GMN created from the mutual information interaction matrix is shown in figure 1d where nodes are denoted M_Vx_ indicating the generalized embedding M of nodes linked to Vx.

**Fig. 1:**
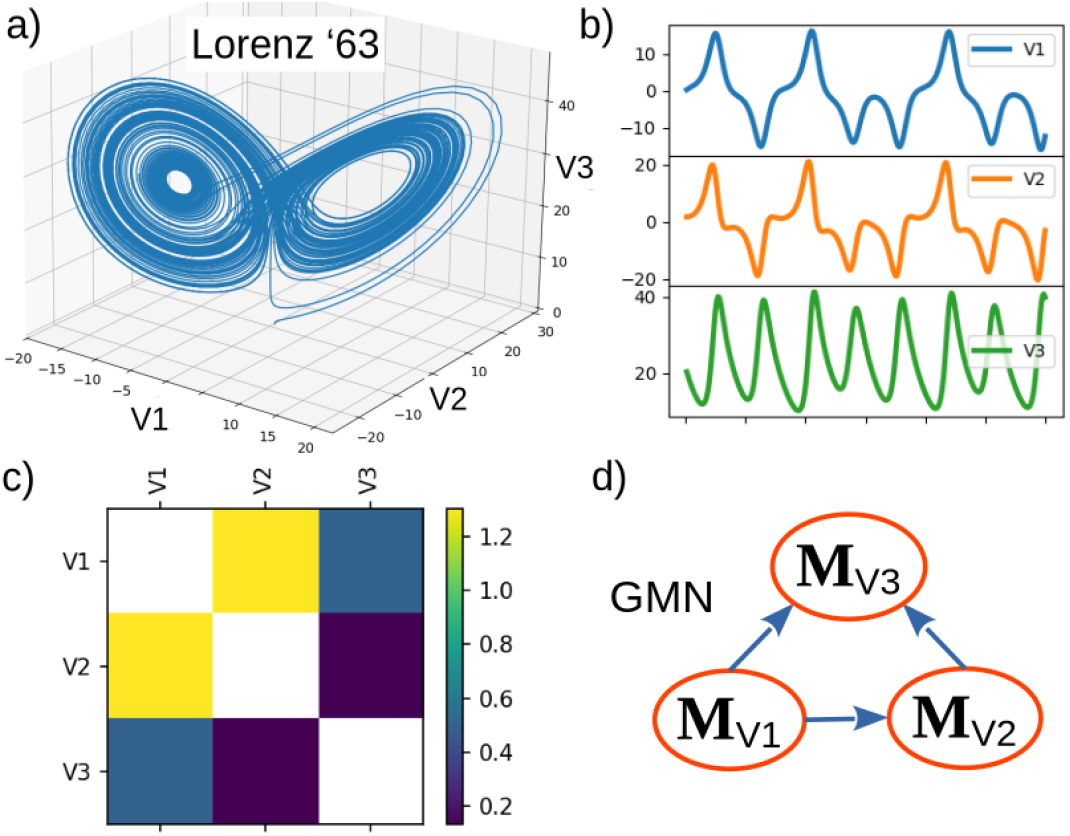
a) The Lorenz’63 system. b) Time series of each variable used to generate mixed-multivariate embeddings for each node of the network. c) Mutual information interaction matrix of the three system variables. d) Three node GMN with node links and embeddings determined from the interaction matrix. Each node is a deterministic, low–dimensional system with a simplex state space generator/prediction function.

Let us consider embedding parameters *E* = 3 and *τ* = − 1. Node M_V1_ has no external inputs and will consist of a 3-dimensional time-delay embedding of variable V1. Node M_V2_ has one input from M_V1_ and will consist of a 3-dimensional time-delay embedding of variable V1 as well as a 3-dimensional time-delay embedding of V2. Such a generalized embedding is termed a *mixed-multivariate embedding*, multivariate indicating more than one observable, mixed referring to a combination of multivariates with time delays. Node M_V3_ receives inputs from M_V1_ and M_V2_ resulting in a generalized embedding with three, 3-dimensional components: a 3-dimensional time-delay embedding of variables V1, V2 and V3.

In generative mode each node produces a prediction of its variable (node M_V1_ predicts V1) at a desired time horizon, here T_p_ = 1 step ahead. The output of each node is recorded and set as the starting point for the next generative time step. This is repeated for a defined number of time steps.

### 1.2 Interaction matrix

A GMN is characterized by the interaction matrix from which the network is discovered allowing one to tailor GMN to address specific aspects of a system. For example, if temporal synchrony is the dynamic of interest correlation can be appropriate. We demonstrate three different interaction functions focusing on specific aspects of each system:

- Mutual information 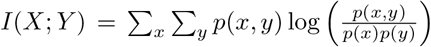 is a metric of shared information. In the Lorenz’63 system we know variables defining the system are causally related, there is no need to seek dynamical relationships focusing on shared dynamics so we use mutual information to recover system interactions based on shared information.
- Nonlinearity *ρ*_Δ_(*x, y*) assesses nonlinear interactions that are *uncorrelated* but nonetheless contribute to shared dynamics: *ρ*_Δ_(*x, y*) = max [*ρ*_*CM*_ (*x, y*), 0]−|*ρ*_*P*_ (*x, y*)| where *ρ*_*P*_ (*x, y*) is linear correlation between observables *x, y* and *ρ*_*CM*_ (*x, y*) the non-linear cross map correlation representing shared dynamics. Specifically, given *x* and *y* a time delay embedding state space of *x* is used to predict observed values of *y* with the simplex predictor [27]. Pearson correlation between predicted and observed values quantify the degree to which states of *x* predict states of *y* indicating shared dynamical information. We use *ρ*_Δ_(*x, y*) in the *Drosophila* example to highlight uncorrelated nonlinear neural interaction, an essential ingredient of neural integration [29, 30].
- Convergent cross mapping (CCM) [18] infers causal relationships ensuring two variables are components of the same dynamical system. In the *Rattus* example we have only partial measurements of whole brain activity and use CCM to identify causally shared dynamics.

Table H1 in supplement H lists other candidate interaction functions.

### 1.3 GMN are universal approximators

Recognition that multilayer neural networks can operate as universal approximators [31] refocused research on deep neural networks. This was followed in control systems where fuzzy controllers were shown to be universal approximators [32], and more recently echo state networks (ESN) have been proven to act as universal approximators [33–35]. An overview of universal approximation theorems is provided in reference [36], a brief review of reservoir computing (RC) and echo state networks with discussion of their connection to random embeddings and Koopman operators is provided in supplement G.

With demonstration that GMN operate on par with ESN in generation of short term synchronized chaotic dynamics and can completely represent complex dynamics, we hypothesized GMN are universal approximators. A proof is provided in supplement B placing GMN in the class of universal approximators, however, in contrast to neural and echo state networks one that is entirely observable and explainable.

## 2 Results

### 2.1 Chaotic dynamics: Lorenz ‘63

The Lorenz’63 atmospheric convection model defines a 3–dimensional manifold according to 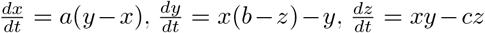 and with parameters a=10, b=28, c=8/3 generates chaotic dynamics. We generate a test set of V1 = x, V2 = y, V3 = z of 4000 points starting at time 10.0 with Δt = 0.02 (figure 1a. With the 3 node GMN of figure 1d using *E* = 3, *τ* = − 5 from a grid search (supplement C) we compare short and long term GMN dynamics with echo state networks consisting of 1000, 2000 and 3000 reservoir nodes. Implementation details are described in Methods.

Both generators are trained on the first 2000 points of data with free-running generation starting at Time = 50 (index 2001) with results shown in figure 2a and 2b where the 1000 node ESN achieves good generative accuracy for approximately 2 seconds. Small improvements can be made with a larger reservoir ESN, but 1000 nodes illustrates reasonable synchronization with the true dynamics. The GMN achieves a comparable but longer period of approximately 2¾ seconds demonstrating GMN is capable of short term accuracy in the generation of chaotic dynamics, but at a remarkably reduced number of free parameters/dimensions in relation to ESN.

**Fig. 2:**
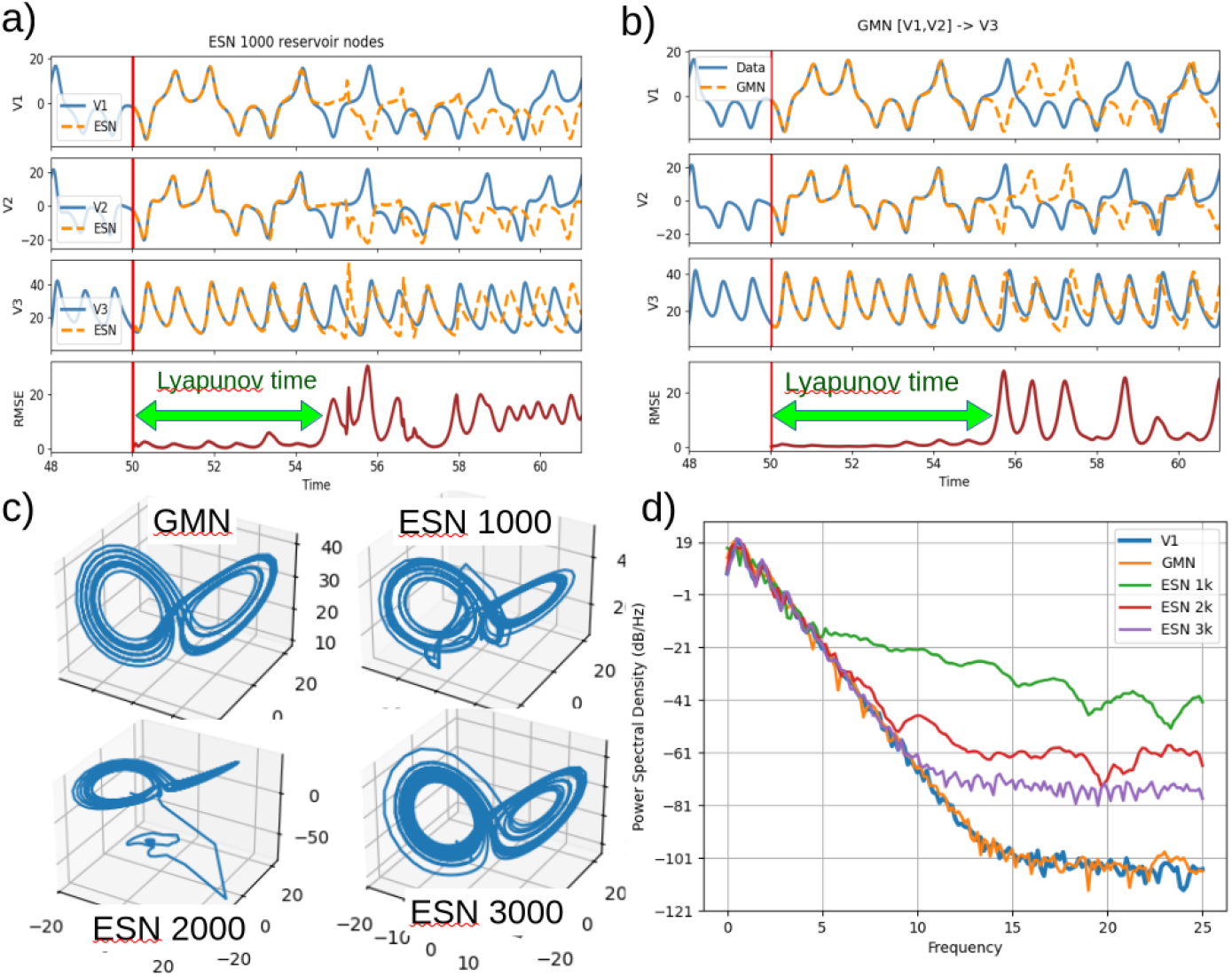
Generated dynamics of the Lorenz’63 system by the 3 node GMN and ESN with 1000, 2000 and 3000 nodes. a) Short term generation of the 1000 node ESN. b) Short term generation of the 3 node GMN. Generative mode begins at Time = 50. c) Long term generation of GMN and ESN with 1000, 2000 and 3000 nodes. d) Power spectrum of long term generated dynamics.

Next we examine long term generation by training both GMN and ESN on the first 2000 points then generating 1000 points as shown in figure 2c. The 3 node GMN generates dynamical behavior consistent with the known dynamics while the ESN requires 3000 reservoir nodes to achieve similar results. To compare properties of the generated time series we calculated the power spectrum of the generated time series indicating the ESN creates unnatural high frequency components. The power of these components decrease with reservoir size but still depart significantly from the spectrum of GMN which is closest to the original Lorenz dynamics (figure 2d).

### 2.2 Drosophila melanogaster

Next we compare GMN and ESN on neural recordings of a fly expressing the calcium indicator GCaMP6f as a measure of neuronal activity. The fly was imaged walking on a freely rotating Styrofoam ball recording forward (FWD) and left/right turning speed (Left Right) as shown in figure 3a. Neural activity was spatially segmented by independent component analysis (ICA) across the brain yielding 80 component brain regions and corresponding time series for a total of 82 observables. Details are provided in reference [37]. The network is created from an interaction matrix based on nonlinearity *ρ*_Δ_(*x, y*) described above. Starting from the forward motion observable (FWD) the network is grown according to the interaction/adjacency matrix as described above and in Methods resulting in a 67 node network shown in figure 3c.

**Fig. 3:**
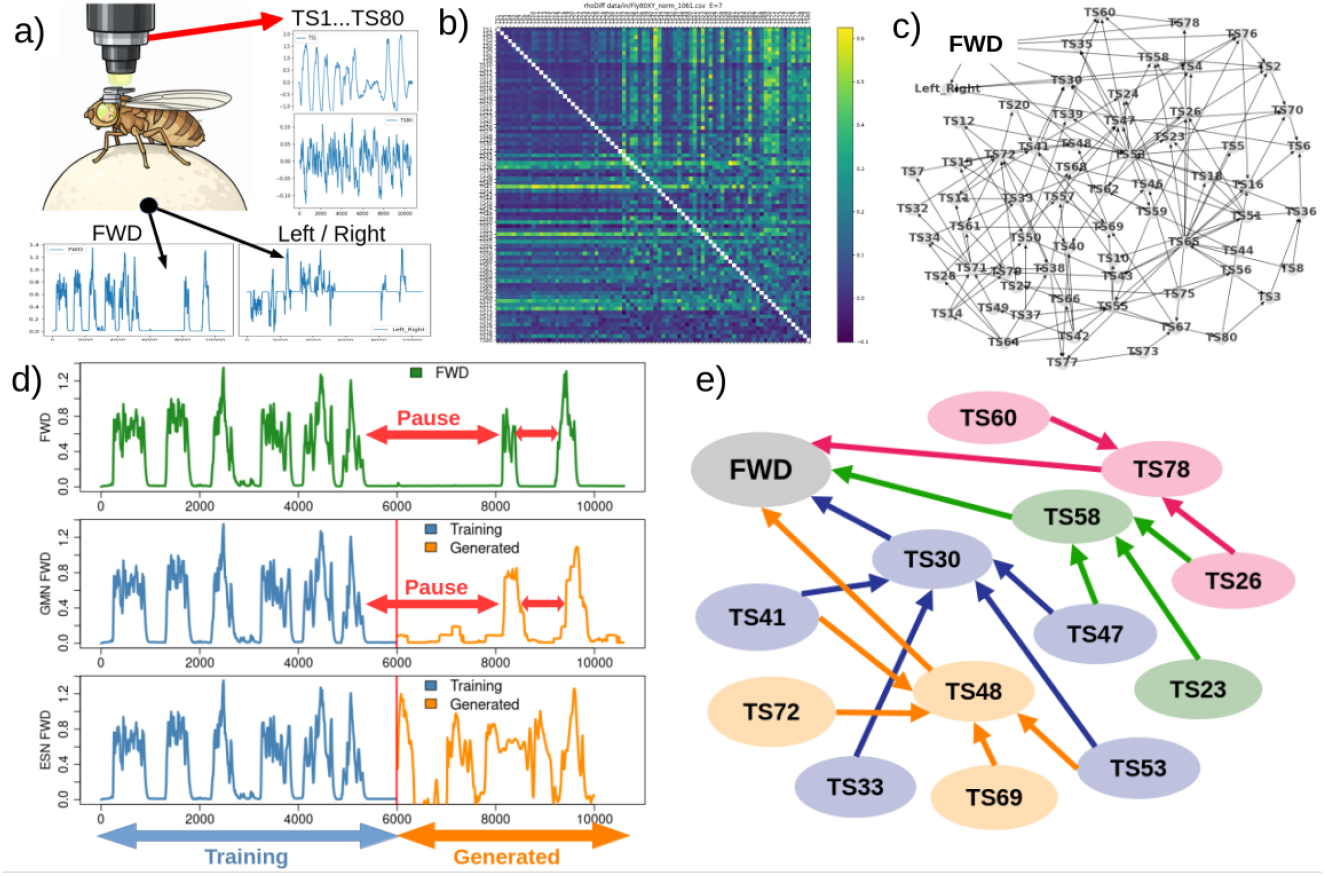
a) Neural and behavioral measurements of *Drosophila* walking on a Styrofoam ball. b) Nonlinear interaction matrix used to create the GMN network. c) GMN network. d) Comparison of forward motion (FWD) observations (top), GMN generated dynamics (middle) and 3000 node ESN generated dynamics (bottom). Cumulative absolute error (CAE) of ESN with respect to FWD is 185.4, GMN 57.4. e) Terminal GMN layers informing FWD.

Comparison of GMN and ESN results are shown in figure 3d where training data span index 1 - 6000, generative dynamics start at index 6001. GMN is run with *E* = 7, *τ* = − 8 for all nodes with *E, τ* selected by a grid search for optimal FWD predictability. Additional generative time series are shown in supplement F.

The training set exhibits six bouts of forward motion separated by small time intervals and does not include the protracted pause in forward motion between the sixth and seventh forward movements. Remarkably, GMN generated dynamics reproduce this pause behavior demonstrating GMN can produce realistic behaviors that are not explicitly present in the training set. This is possible from interactions of multivariate, multiscale generalized embeddings that capture system dynamics across scales from multiple variables as well as delays thereof. Removal of either of these features, node interactions or multiscale embedding precludes this ability as verified with autonomous generation of dynamics from a univariate time delay embedding (no network interactions) and from multivariate embeddings with *E* = 1 (no multiscale dynamics), neither of which generate the observed pause (supplement E). We therefore infer GMN are capable of learning the complete dynamics of a system as long as the observations are sufficiently informative, the network structure sufficiently connected, and time delays appropriate. This inference is supported by the GMN universal approximation theorem asserting GMN are capable of predicting any global network state as long as the library of states are sufficiently dense (supplement B).

In contrast, ESN generation of *Drosophila* forward motion fails to reproduce characteristics of the observed time series in the withheld data. The ESN generated time series start immediately with a fast bout then producing multiple bouts in a quick succession rather than a long pause and two bouts in the withheld data. A direct comparison of GMN & ESN generation is shown in figure 3d where the ESN is observed to generate unrealistic negative forward motions while GMN generated dynamics better approximate the withheld data. It should be noted in principle ESN results can be improved perhaps with larger reservoirs, deep ESN, or next generation ESN (adding explicit nonlinear outputs or internal states) [38], however, one is then engaged in a protracted optimization exercise with a reservoir that may be finely tuned to the specific behavior [39], and, one has still not recovered mechanistic insight derived from observables.

Regarding explainability figure 3e shows the last two GMN layers informing forward motion where nodes TS30, TS48, TS58, TS78 directly inform FWD dynamics with each of these informed by multiple, integrated nodes in the second layer. Specifically, TS30 is located in the Optic Lobe (OL), a key region of sensory integration, TS48 the Mushroom Body (MB) effecting sensory integration, TS58 the Central Complex (CX) engaged in motor control and spatial navigation, and TS78 in the Lateral Complex (LX) connecting sensory inputs with motor functions. Second-tier regions informing these integrative functions include TS26, TS33, TS41, TS53, TS60, TS72, all located in the Optic Lobe, TS23 of the Gnathal Ganglia (GNG) associated with feeding and TS69 in the Superior Neuropil (SNP) integrating sensory and motor processing.

Thus GMN has plausibly identified neural pathways associated with integration of sensory and motor functions of forward movement. Generally, identification of regions driving observed behavior sets the stage for experimental manipulation to validate the role of upstream nodes, an effort beyond the current scope but currently underway.

### 2.3 Rattus norvegicus

GMN and ESN were applied to neural and behavioral data of a freely-moving rat in a fork-maze performing a spatial task. In this task the rat was rewarded only if it alternated visits from a central arm to the left and right arms (e.g. center - left - center - right - center …). The animal was implanted with a combination of four wire electrodes targeting the CA1 region of the hippocampus and 128 electrode flexible polymer devices targeting the dorso-medial prefrontal cortex (mPFC) and orbitofrontal cortex (OFC) regions. The dataset chosen for this analysis was from a later stage when the animal had mastered the task and includes 468 units (putative single neurons) recorded simultaneously from these three regions. Animal position was recorded by an overhead camera yielding X and Y coordinates, with the track oriented such that X position relates to which track segment the animal is in while Y position relates to where in that segment the animal is located as shown in figure 4a.

**Fig. 4:**
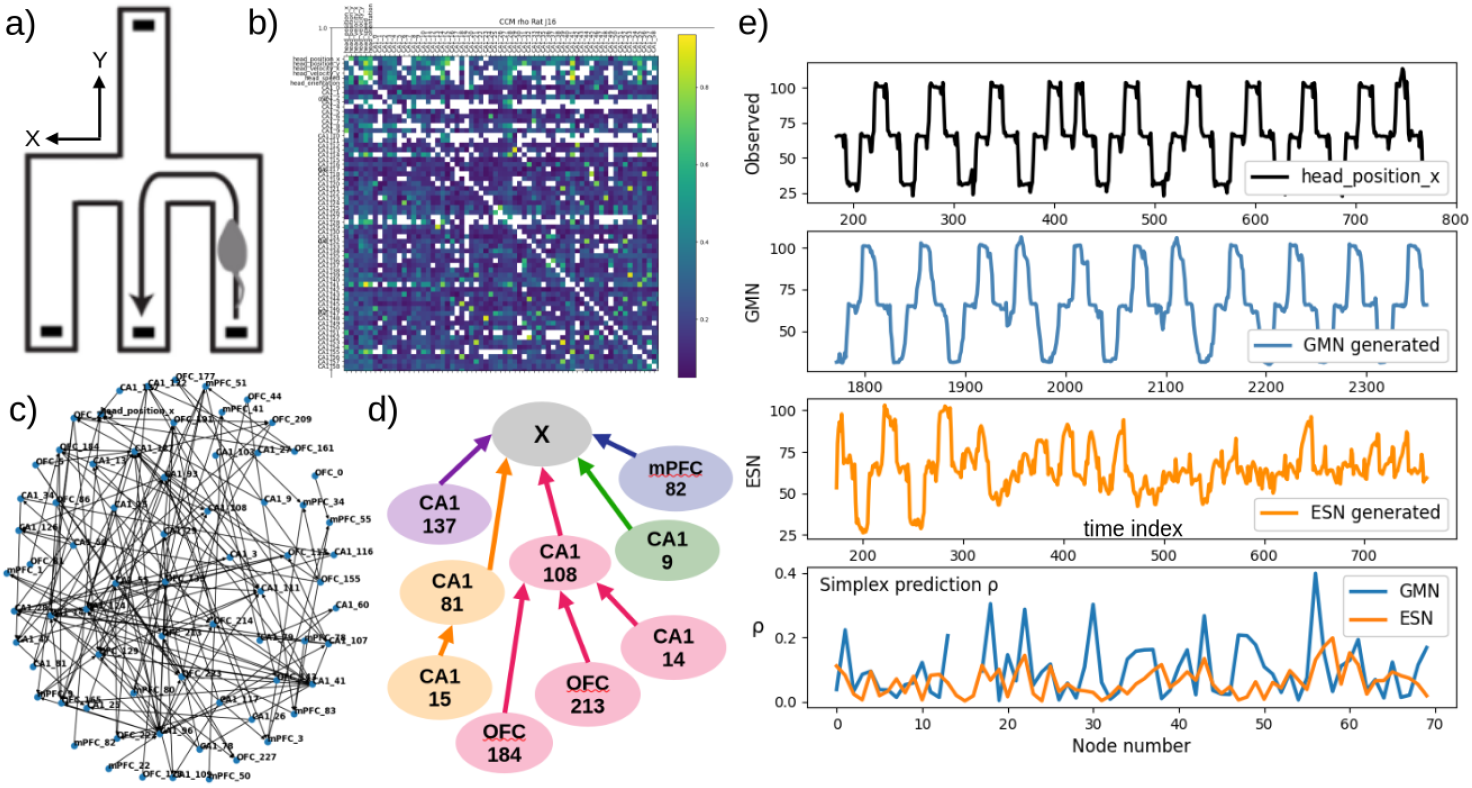
a) Schematic of maze with reward wells at the terminus of each segment. b) Convergent cross mapping (CCM) interaction matrix of neural and behavioral data. c) GMN created from CCM interaction matrix. d) Terminal GMN layers informing position X. e) Comparison of observed position X (top), GMN generated positions, ESN generated positions and simplex prediction skill of observed variables from time delay embeddings of GMN and ESN generated time series.

The interaction matrix is computed from convergent cross mapping (CCM) as shown in figure 4b . This matrix yields a network focusing on causal interactions ensuring linked observables are contributing to node dynamics. Empty (white) cells correspond to CCM interactions that failed to converge. The last two layers of the network informing position X are shown in figure 4d revealing hippocampal (CA1) domination. The identified regions are principled candidates for experimental manipulation to explore the importance of these neurons in the representation of animal position during the task.

Comparison of GMN generated position X with a 50,000 node ESN is shown in figure 4e. It is evident GMN generated dynamics are representative of the actual dynamics while ESN values lack essential characteristics of the observed dynamics after a short period. To assess the fidelity of generated compared to observed dynamics we use the GMN generated time series at each node to create a time-delayed embedding of the synthetic time series from which simplex predictions are made of the observed data. The bottom panel of 4d) shows the Pearson correlation between predicted and observed data demonstrating GMN generated dynamics generally outperform ESN generated dynamics and hence are a better description of the system than ESN generated embedded data.

### 2.4 Crossformer

Deep neural networks continue to undergo significant advancement as machine learning leverages the *transformer* architecture where positional encodings encapsulate long range correlations and where recurrence and convolution are replaced with stacked self-attention and feed-forward layers facilitating parallelizable sequence modeling [40]. Transformer applications have recently focused on multivariate time series [41] of which the crossformer is a top benchmark performer by adding cross-dimension dependency (relationships across variables) to the usual cross-time dependency [42].

Here we compare GMN with crossformer on the standard Electricity Transformer Dataset [43] with results shown in figure 5 where we find crossformer predictions are considerably less accurate than GMN. These differences likely reflect that on long horizons transformers rely mainly on trends/periodicity rather than characterization of temporal dynamics, and are prone to overfitting noisy/aperiodic data, but at high computational cost [44]. Further, transformer architectures are specifically tailored to benchmarks and it has been shown a significant portion of benchmark success derives from Z-score normalization [45]. Indeed, in the comparison of figure 5 one can note the crossformer variables have been transformed from observation space to normalized values, while GMN values are mappings to the true observed variables.

**Fig. 5:**
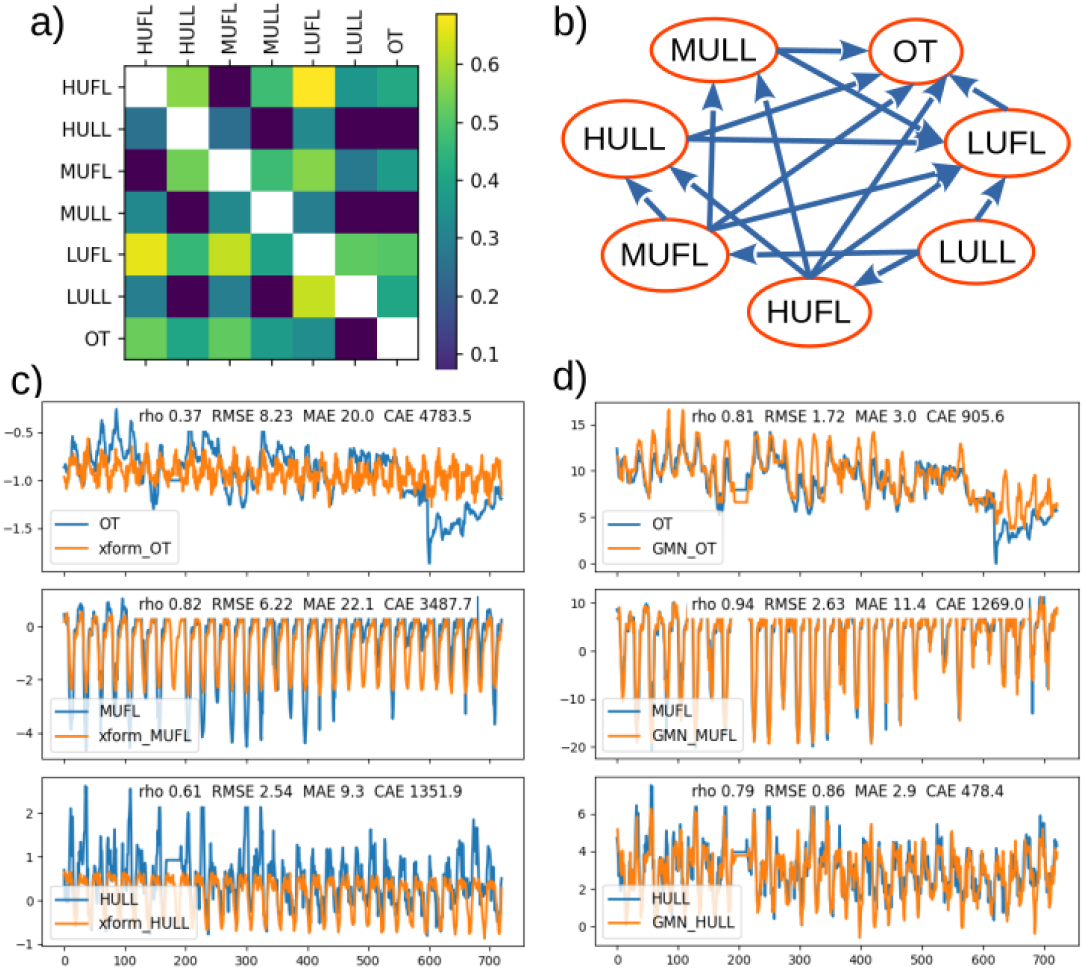
a) Convergent cross mapping (CCM) interaction matrix of electrical transformer data. b) GMN created from CCM interaction matrix. c) Crossfomer predictions of variables OT (Oil Temperature), MUFL (Middle UseFul Load), HULL (High UseLess Load). d) GMN predictions of OT, MUFL, HULL. Error metrics are Pearson correlation between predictions and observations (rho), Root mean square error (RMSE), Maximum absolute error (MAE) and Cumulative absolute error (CAE).

## 3 Discussion

A recurring theme in neuroscience and other complex systems is that high-dimensional observations often evolve on a much lower-dimensional structure [1]. In neuroscience for example, population activity can frequently be described by a relatively small number of variables, yet those variables are typically modeled in latent spaces defined by matrix factorization, principal components, or deep learning representations. While latent representations are often powerful for compression and prediction they can obscure contributions of individual observed variables, their interactions, and the timescales over which those interactions unfold [16, 17].

Generative manifold networks provide an alternative formulation. Rather than representing system dynamics in a single latent space, GMN models a system as a network of interacting, low-dimensional manifolds anchored directly to observables. Each node consists of a generalized embedding of measured variables and their delays, with network structure determined by an interaction criterion chosen to express a property of interest such as causality, shared information or nonlinearity. In this sense, GMN combines three desirable features that are often difficult to obtain simultaneously: low-dimensional representation, explicit network structure, and direct interpretability in observation space. Furthermore the quality of the model can be quantified by its ability to predict system behavior with out of sample data and/or simulate (generate) the system dynamics and evaluate the realism of such simulation, both objective metrics of success. Because the model is data-driven and does not rely on latent or randomly initialized variables it yields a representation that is directly testable against experiment while remaining expressive enough to serve as a universal approximator limited only by data and suitable network representation.

The empirical results suggest this architecture captures aspects of neural and behavioral dynamics that are difficult to capture with more conventional approaches. In the Drosophila example, GMN reproduced a prolonged pause in forward motion that was not explicitly represented as a repeated motif in the training segment, while echo state networks generated less realistic trajectories and required substantially larger state dimension. In the rat dataset, GMN generated dynamics that better preserved the structure of the observed animal positions in relation to neural signals and identified interpretable upstream contributors dominated by hippocampal inputs consistent with known roles of CA1 place cell neurons. Together, these results support the view that GMN can recover system-level dynamics while retaining mechanistic links to measured variables, an especially important property in neuroscience where models are most useful when they can suggest candidate circuits, regions, or populations for perturbation and validation.

Two features appear to be especially important for this performance: multiscale temporal structure and network interactions. First, the Drosophila analyses show that removing delay coordinates prevents GMN from reproducing key behavioral structure. This finding implies that relevant state information is distributed across timescales and may not be fully present in instantaneous observations alone. In this respect, GMN leverages the logic of generalized state-space reconstruction: delays act as surrogates for unobserved or slow variables and allow local manifolds to represent dynamics that are only partially measured [22]. Second, network structure is also essential. A single manifold, when run autonomously, can collapse to an unrealistic limit cycle, whereas a network of interacting manifolds sustains richer free-running dynamics. These observations argue that realistic generation in complex neural-behavioral systems requires both multiscale embeddings and explicit interactions among components.

For neuroscience, this point is conceptually important. Neural systems are neither purely feedforward nor adequately described by a single global trajectory detached from anatomy and circuit organization. GMN instead offers a middle ground between highly abstract latent models and detailed mechanistic simulations. It preserves node-level observability while allowing distributed interactions across regions, cells, or behavioral variables. The resulting networks are therefore not merely predictive; they are also hypothesis generating. In the fly and rat examples the terminal layers of the GMN identify candidate upstream regions associated with behavioral output. Such structure can, in principle, guide lesion, stimulation, or recording experiments aimed at testing whether the inferred pathways are causally involved in the behavior of interest.

Although ablation studies are a popular way to try and identify intervariable dependencies, they are founded on separability of component interactions and therefore GMN is not ideal for virtual lesion experiments that would delete variables/observables and test the resulting consequences. The reason is the inherent nonseparability of nonlinear systems and Takens-style reconstruction: each observed time series partially contains information about other interacting variables, including unobserved ones. As a result, removing one variable from an embedding does not fully remove its influence since the influence is already distributed across the remaining observables, all of which were measured in its presence [18]. Nonetheless, one can test the reconstruction quality of a network topology by severing putative causal connectivity of the simulation as shown in Supplement D. The effectiveness of such an approach is likely constrained by the limits of observability at each node [46, 47]

The present implementation also clarifies several boundaries of the approach. GMN currently generates dynamics using a manifold library constructed from the training data, and free-running trajectories are produced by iteratively updating within the learned manifold. Thus, while the model can generate rich autonomous dynamics the underlying manifold itself is static during generation. An important next step will be to allow online updating of the manifold or adaptive generator rules enabling continual learning in settings where the system changes over time. This will be particularly relevant for neural data acquired across learning, context shifts, or nonstationary internal states.

A second practical issue is model construction. Although GMN is conceptually simple, performance depends on choosing an interaction function appropriate to the scientific question and embedding parameters that capture relevant timescales. This dependence should be viewed less as a weakness than as a strength of the framework: it allows the investigator to tailor the model toward causal interactions, nonlinear integration, synchronized dynamics, or other properties of interest. At the same time, principled model selection remains important, especially for large neural datasets in which multiple network topologies and embeddings may perform similarly well. More broadly, we find that viable GMN configurations are often relatively easy to obtain which may reflect a general complex system principle that many different parameter settings yield similarly good solutions as described by sloppy parameter sensitivity and flat minima as well known in the mathematical literature [48–52].

The computational properties of GMN are also encouraging. Although the method scales linearly with node number and node dimension it can leverage GPU-accelerated empirical dynamic modeling backends and should remain practical for large systems [53]. This opens the possibility of using GMN not only as an analysis tool for neural recordings, but also as an interpretable surrogate model in coupled or hybrid settings, where one seeks a computationally efficient representation of a complex dynamical subsystem without sacrificing observability.

More broadly, GMN suggests a different way to think about low-dimensional structure in neuroscience. Instead of asking how to compress neural data into a single latent manifold, GMN asks how low-dimensional manifolds attached to observed variables interact to generate collective dynamics. Such a shift is important because it reframes dimensionality reduction from a primarily descriptive exercise into a generative and experimentally actionable one. In systems where explanation and experimental verification matter as much as prediction, this may be a useful organizing principle[54].

In summary, GMN provides an interpretable, data-driven framework for prediction and free-running simulation of complex dynamics that integrates multiscale state-space reconstruction with explicit network structure. The results here suggest that this combination can capture behaviorally relevant neural dynamics with substantially lower dimensionality than standard reservoir-based methods while preserving direct links to measured observables. For neuroscience, the major opportunity is not only improved forecasting, but the prospect of models that point back to circuits, timescales, and interactions that can be tested experimentally.

Leveraging the significant dimensional reduction and transparent, explainable basis of GMN, it should find broad applicability as a surrogate model in coupled/hybrid model applications where surrogate (embedded) models of complex dynamics replace computationally intensive numerical model components [55].

## 4 Methods

Generative manifold networks are conceptually and operationally simple consisting of a network of interacting dynamical systems generating time series at each node of the network. GMN is data–driven with the network discovered from an interaction/adjacency matrix, and node generation by a multi-input single-output (MISO) function. GMN can therefore be completely configured with the choice of interaction and generator functions.

The network interaction function can be any comparative or discrimination function such as convergent cross mapping, nonlinearity or correlation. Several candidate interaction functions are detailed in supplement H. In the present work nodes consist of generalized embeddings [22] with a simplex state space generator [27] characterized with two parameters, the embedding dimension *E* and time delay *τ* . However, each node can have a distinct generator with node-specific parameters. Values of *E* and *τ* can be determined by assessing optimal simplex predictive skill of the target variable over ranges of *E* and *τ* .

Results presented here are computed with the python GMN package `dimx` [56] and EDM package `pyEDM` [57]. Code and data to reproduce the results are available at GMN_ESN_Examples.

### 4.1 Algorithm

Given an N row by M column observation matrix O_NM_ where M corresponds to observables and N to time series instances an MxM interaction matrix I_MM_ is computed. The interaction function can be any suitable metric appropriate to classification of the dynamics such as CCM, mutual information, nonlinearity or correlation. A list of candidate interaction functions is provided in supplement H. With the interaction matrix I_MM_ the associated GMN is implemented in two steps:

1. Create Network
  □ Starting at a desired target node selected from the M observables, add up to N_D_ driver nodes for the target according to the interaction/adjacency matrix while disallowing network cycles. N_D_ can be determined as a dimensional estimate of the target behavior, or, as a node–specific dimensional estimate with the `pyEDM` function `EmbedDimension` where the node observable is time delay embedded into a state space and simplex predictive skill assessed as a function of embedding dimension. The embedding dimension with optimal predictability is selected as the number of node inputs N_D_.
  □ Repeat at each connected node until no more connections.
2. Generate Dynamics
  □ The GMN state space manifold 𝕄 consisting of the ensemble of node embeddings is created from a specified time range of observations.
  □ A forward prediction is made at each node from the manifold 𝕄. The set of predictions from all nodes constitutes the network output at that time step, the one-step ahead manifold 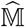.
  □ Forward predictions are repeated for a defined number of steps. At each step the previous network output 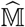 defines the state from which the next time step prediction will be made.

The GMN algorithm is detailed in supplement A. Proof that GMN are universal is provided in supplement B.

## Declarations

### Data availability

Data are available at GMN_ESN_Examples.

### Code availability

Code is available at GMN_ESN_Examples.

### Author contribution

GMP conceived and designed GMN. JP and CS designed and implemented GMN. GMP, JP and CS applied and evaluated GMN, and contributed to the manuscript. JP generated the GMN universality proof, AS and SS revised the proof. JG generated the Rat data which was processed by SYT under the supervision of LMF.

## Acknowledgments

The present work was supported by the Okinawa Institute for Science and Technology internal core funding, a grant from the W.M. Keck Foundation. We thank Sophie Aimon for providing us with her original lightfield microscopy data of walking *Drosophila melanogaster* from her paper[37]. We thank Seann Wang for critical reading of the manuscript.

The funders had no role in study design, data collection and analysis, decision to publish, or preparation of the manuscript.

## Supplement A GMN Algorithm

### A.0.1 Create Network

Given an interaction matrix I_MM_, target observation node, node functions and number of node drivers (inputs) create the network graph G, see algorithm 1.

#### Algorithm 1

Create Network

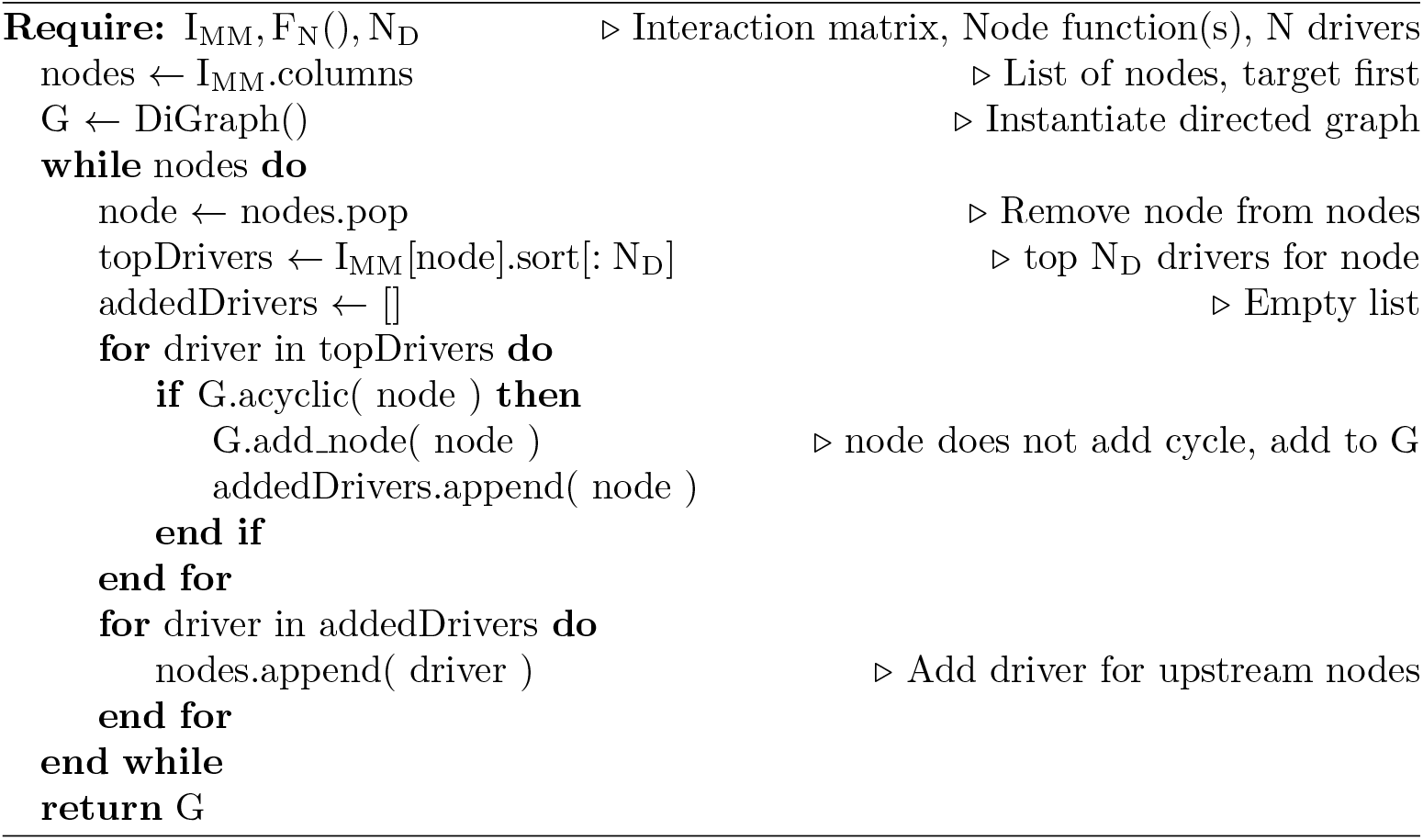

### A.0.2 Generate

Given a GMN graph G, observation matrix O_NM_ and training indices, generate dynamics, see algorithm 2.

#### Algorithm 2

Generate

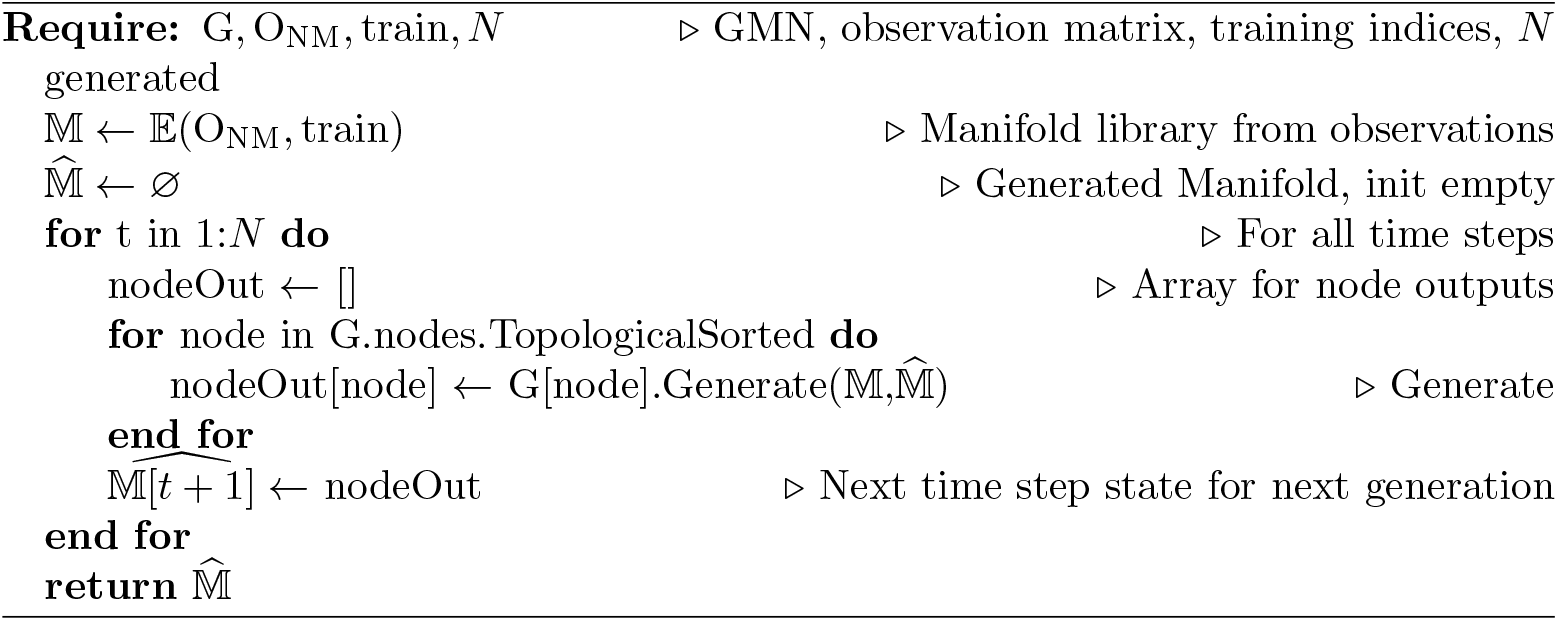

## Supplement B Generative manifold network universal hypothesis and proof

Here we state a universality theorem for a global network update operator on a joint reconstructed attractor in terms of the adjacency matrix and mixed embedding operator. The proof is based on a supremum norm of GMN error over the attractor.

### B.1 Definitions

#### B.1.1 Networked dynamical systems

Let an underlying system evolve on a compact invariant set, the attractor 𝒜 inside a smooth manifold ℳ:

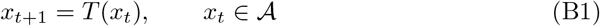

with *T* : 𝒜 → 𝒜 continuous. Let node observables be

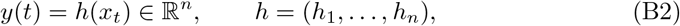

where each *h*_*i*_ : 𝒜 → ℝ is smooth, meaning it has extension to a neighborhood of 𝒜 in ℳ, in which it is a smooth map.

#### B.1.2 Adjacency matrix and embedding operator

GMN is defined by a directed adjacency matrix of rank *n*

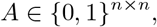

where *A*_*ij*_ = 1 if and only if *j* ∈ *P*_*i*_(*A*), where *P*_*i*_(*A*) is the set of parent nodes for *i*. We assume self-information is always allowed corresponding to *A*_*ii*_ = 1.

Let *τ >* 0 be fixed. Define *θ* as mixed embedding parameters that specify for each ordered pair (*i, j*) with *A*_*ij*_ = 1 a finite set of delays (lags)

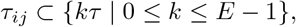

where *E* equals the embedding dimension from Takens theorem. Define node *i*’s mixed delay embedding ℰ as the map defined along system trajectories with *P*_*i*_(*A*) := *j* : *A*_*ij*_ = 1 ∪ *i* defined as node *i*’s parent set. Define

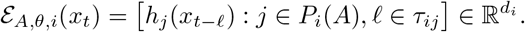

Let *E*_*ij*_ = dim ℰ _*A,θ,i*_(*x*_*t*_). Put 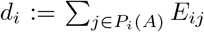 the total embedding dimensions at node *i*.

Now form the joint GMN embedding state representing the GMN global state:

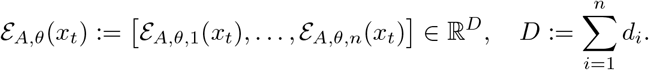

Define the joint reconstructed attractor

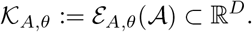

𝒦_*A,θ*_ is compact because it is the continuous image of a compact set.

#### B.1.3 Global network update operator on the joint attractor

On 𝒦_*A,θ*_, define the *true* global update operator

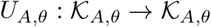

by conjugacy (equivalence under homeomorphism):

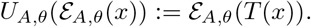

and since *T* (*x*) is the one step ahead evolution operator on *x* this reflects advancing the underlying system by one step and updating the embedding.

### B.2 The GMN hypothesis

A GMN with simplex generators is specified by:

- Adjacency *A*
- Embedding parameters *θ*
- A finite library of embedding states *z*_*i*_ and target values *v*_*i*_ for each node *i* :

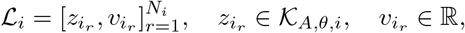
- A simplex generator at each node *i* : for input *z*_*i*_, pick its *k* nearest library points 𝒩_*k*_(*z*_*i*_) and output the convex combination

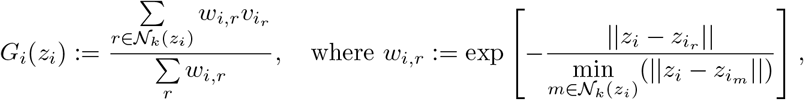

where the norm is taken to be the standard Euclidean norm. The GMN global one-step output map is *G* : 𝒦_*A,θ*_ → ℝ^*n*^ defined as

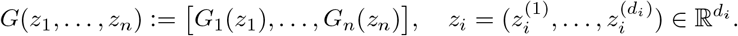

For a vector 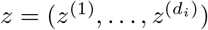 let 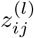 be *j*-th coordinate of *z*^(*i*)^ with given time lag *l*. Now put

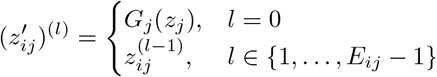

which means 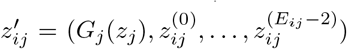. Now using notations above define the GMN global update operator as

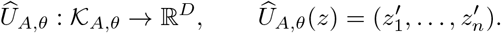

The GMN hypothesis class is the union over all admissible (*A, θ*) and all libraries/weights:

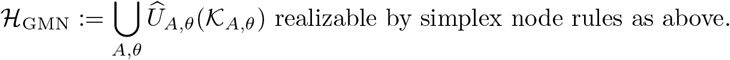

### B.3 GMN universality theorem

*Theorem for universality of GMN with simplex generators on the global update operator*.

Let both {*h*_*i*_} and {*τ*_*ij*_} be generic in the sense of smooth manifolds and let the causal adjacency matrix *A* sufficiently represent the global dynamics through {*h*_*i*_}. Then

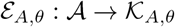

is an injective homeomorphism, which can be extended to diffeomorphism onto its image.

Then:

For any *ε* > 0, there exists a simplex GMN in ℋ_GMN_ using (*A, θ*) such that the induced global update operator Û_*A,θ*_ satisfies

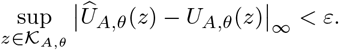

In particular, the family of all GMN global operators, parameterized by the library size, embedding dimensions, and neighborhood size *k*, is dense in the space of continuous mappings *C*(𝒦_*A,θ*_, 𝒦_*A,θ*_) with respect to the supremum topology. Equivalently, for any true continuous operator *U*_*A,θ*_ and any *ε >* 0, there exists a GMN operator *Û*_*A,θ*_ ∈ ℱ_*GMN*_ such that sup_*z*_ ∥*Û*_*A,θ*_(*z*) − *U*_*A,θ*_(*z*)∥_∞_ *< ε*.

### B.4 Sufficient conditions for node embedding

Let the total system variables be *V* = *V*_1_ ∪ *V*_2_, where *V*_1_ are the observable variables and *V*_2_ are unobserved. The node embedding assumption holds generically if:

- The unobserved variables *V*_2_ are dynamically coupled to the observables *V*_1_ (there are no invariant sub-manifolds completely decoupled from the measurement functions);
- embedding dimension of the local state vector *z*_*i*_, defined as 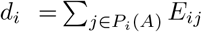, satisfies *d*_*i*_ *>* 2*d*, where d is the box-counting dimension of the global attractor 𝒜;
- The measurement functions and time delays are generic (avoiding underlying symmetries or periodicities of the attractor).

### B.5 GMN universality proof

#### B.5.1 The global update operator is continuous on 𝒦_***A***,***θ***_

As shown above, since the global update operator applies the time evolution operator *U*_*A,θ*_(ℰ_*A,θ*_(*x*)) = ℰ_*A,θ*_(*T* (*x*)) and since *T* and ℰ_*A,θ*_ are continuous, *U*_*A,θ*_ is continuous on 𝒦_*A,θ*_. The set 𝒦_*A,θ*_ is compact, so *U*_*A,θ*_ is uniformly continuous. Hence, for any *ε >* 0 there exists *δ >* 0 such that

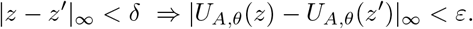

#### B.5.2 Reduce to approximating the update by simplex

We can express the joint update as a deterministic shift-and-append:

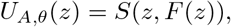

where the shift operator *S* : ℝ^*d*^ *×*ℝ → ℝ^*d*^ is defined as *S*(*z, v*) = (*v, z*_1_, …, *z*_*d*−1_) fora state vector *z* = (*z*_1_, …, *z*_*d*_) and a generated scalar *v* = *G*(*z*). Endowing the product space with the supremum norm || (*z, v*) || _∞_ = max(||*z*||_∞_, | *v*|), it follows directly that for any (*z, v*) and 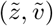:

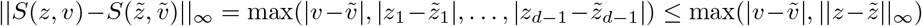

Thus, *S* is globally 1-Lipschitz. Consequently, the continuity of the full update operator *Û*(*z*) = *S*(*z, G*(*z*)) is strictly determined by the continuity of the generator function *G*(*z*).

If one can build a GMN output map *G* such that

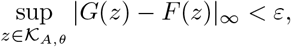

then automatically

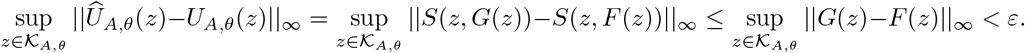

So it suffices to approximate *F* uniformly.

#### B.5.3 Construct nodewise target functions on nodewise attractors

As above we assume the mixed embedding at node *i* is rich enough that the next value *y*_*i*_(*t* + 1) is a continuous function of *z*_*i*_ on the attractor. A sufficient (strong) condition is the node embedding assumption, that each ℰ_*A,θ,i*_ is itself a homeomorphic embedding of 𝒜 onto *K*_*A,θ,i*_, and require GMN-locality: node *i* can only apply its function to its local embedding *z*_*i*_ ∈ 𝒦_*A,θ,i*_.

For each node *i*, define its nodewise embedded attractor

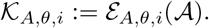

and since the update operator applies the time evolution operator on observables *U*_*A,θ*_(ℰ_*A,θ*_(*x*)) = ℰ_*A,θ*_(*T* (*x*)) there is a mapping to embedding states

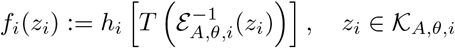

Since each *f*_*i*_ is continuous on compact 𝒦_*A,θ,i*_ it is uniformly continuous over 𝒦_*A,θ,i*_.

#### B.5.4 Approximate each *f*_*i*_ by a simplex node generator

Fix *ε >* 0. For node *i* by uniform continuity of *f*_*i*_ one can choose *δ*_*i*_ *>* 0 such that

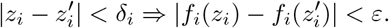

Since 𝒦_*A,θ,i*_ is compact, choose a finite library 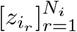 such that for every *z*_*i*_ ∈ 𝒦_*A,θ,I*_ its *k* nearest library points lie within *δ*_*i*_. Set targets 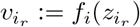. As above the simplex predictor is

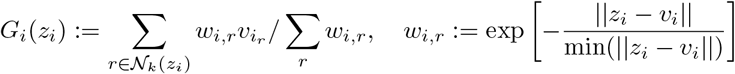

Then for any *z*_*i*_ ∈ *𝒦*_*A,θ,i*_, every neighbor 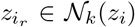 satisfies 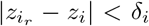 hence 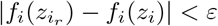. Since the simplex operator is convex in *z*_*i*_

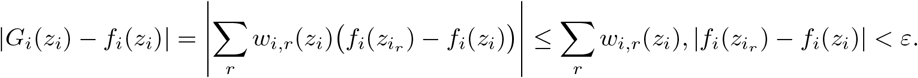

Therefore,

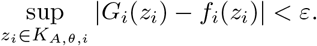

#### B.5.5 Combine nodes for uniform bound on the global output map *F*

On the joint attractor 𝒦_*A,θ*_, each point has the form

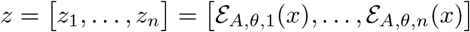

for some *x* ∈ 𝒜. The true next-output is

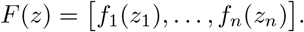

The GMN output map is

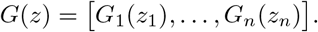

Then, using the sup norm,

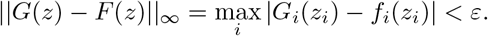

So

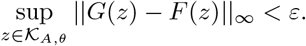

#### B.5.6 Lift to the global update operator

As shown above, since Û_*A,θ*_(*z*) = *S*(*z, G*(*z*)) and *U*_*A,θ*_(*z*) = *S*(*z, F* (*z*)) with *S* 1-Lipschitz in the updated estimate

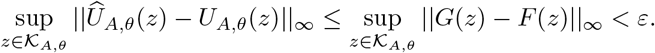

### B.6 Comments

- The adjacency matrix *A* and mixed embedding operator ℰ_*A,θ*_ are part of the model class specified by the data.
- Under the node embedding conditions established in Section B.3 (which guarantee dynamics on 𝒦_*A,θ*_ are Markovian), the map to *z*(*t* + 1) is uniquely determined and deterministic on the mixed embedding manifold, then:

A simplex-based GMN can approximate the entire global network update operator uniformly on the joint attractor to arbitrary precision by taking a sufficiently dense library.

## Supplement C Selection of *E τ* embedding parameters

To select embedding parameters *E,τ* a grid search of simplex model predictability over values *E,τ* can be performed. For example, Figure C1 plots a grid search over *E,τ* for the Lorenz’63 GMN. Enclosed voxels show candidate embedding parameters that produce Pearson correlation (rho) between generated and observed dynamics greater than 0.94.

**Fig. C1:**
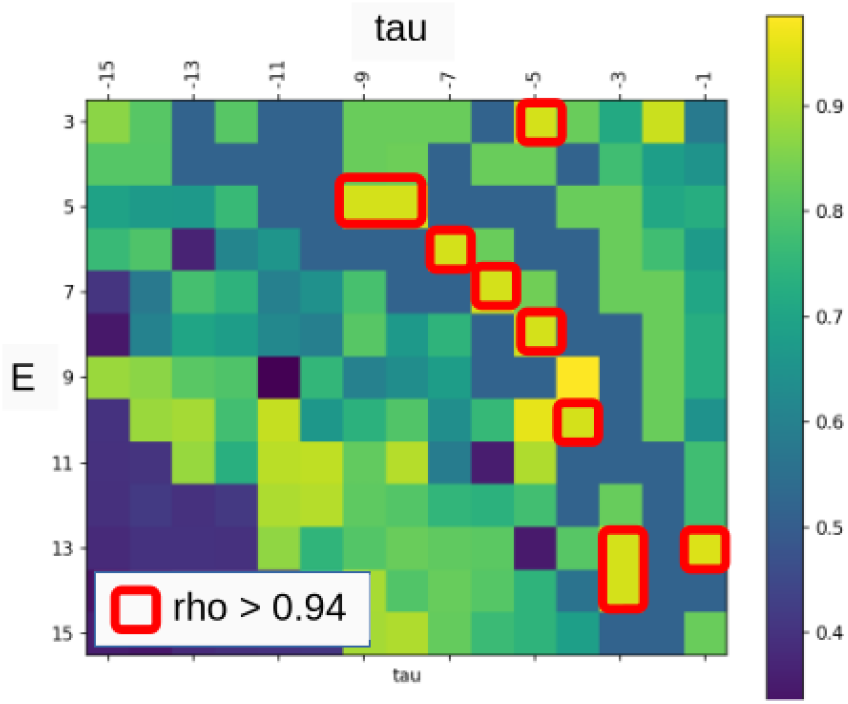
Pearson correlation (rho) between generated and observed dynamics of the Lorenz’63 GMN as a function of embedding parameters *E,τ* .

## Supplement D Network manipulation

Given an interaction function, network discovery proceeds by traversing the adjacency matrix from the target node adding input nodes according to interaction rank while avoiding cycles. Each added node is then recursively explored until no further nodes can be processed. This algorithm explores the interaction structure resulting in a directed graph of information flow. To assess contribution of specific nodes/links, network ablation can be performed.

For example, in figure D2a the network discovered from mutual information of the Lorenz’63 system is shown with its generated dynamics faithfully reproducing the known dynamics. Removal of the V1 to V2 interaction is shown in figure D2b demonstrating loss of crucial dynamical information.

**Fig. D2:**
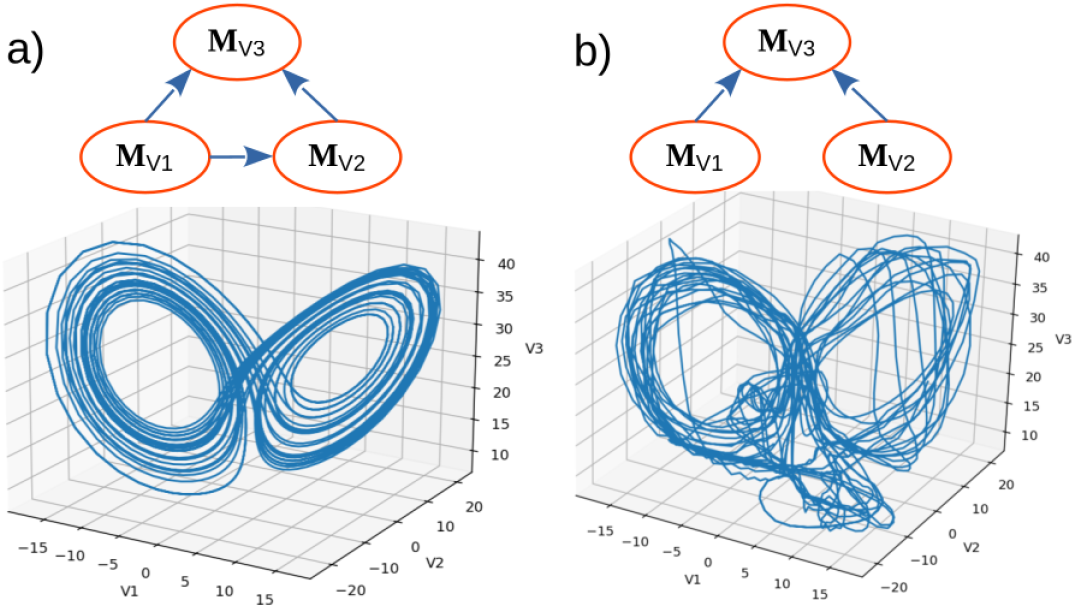
a) A complete network generates full dynamics. b) An incomplete network fails to generate high fidelity dynamics.

## Supplement E GMN multiscale interactions

The original Takens [28] theorem guarantees that under generic conditions (measurements are coupled to all relevant degrees of freedom in the system)a delay embedding of a single variable of a multivariate system allows reconstruction of a manifold that is diffeomorphic to the original system. The delayed vectors can be thought of as surrogates of unobserved variables that potentially act at different timescales. The 2011 generalization of the Takens theorem [22] validated mixing of delays and variables into mixed-multivariate embeddings facilitating multiple timescales within the combinations. In fact the multiple timescales can be incorporated as non uniform delays as shown in figure E3 panel E where delays have a magnitude of 10 in the horizontal axis and a 3X larger delay of 30 in the vertical axis. This differs from the original Takens application where a single dominant timescale was taken into account as shown in figure E3 A, B, where each axis has delays that are required to be integer multiples of the delay magnitude tau which is 30 in this case.

In the *Drosophila* example we found that both network interaction and multi-scale dynamics were required to generate realistic dynamics of fly forward motion. Here we demonstrate those results corroborating that both network structure and multiscale dynamics are essential when modeling complex dynamics from incomplete observations.

The top six panels of figure E4 compare observed time series (left column) with time series generated by GMN lacking multiscale information. Here, all GMN nodes have an embedding dimension E=1 preventing any multiscale temporal information in a time delay embedding. In all cases the generated dynamics fail to reproduce the observed dynamics.

The bottom panel of figure E4 shows fly FWD motion generated from a single manifold (no network) with embedding dimension E=7 and embedding delay tau=-1. Here we find the single manifold provides poor fidelity to the observed behavior eventually becoming trapped in a limit cycle.

**Fig. E3:**
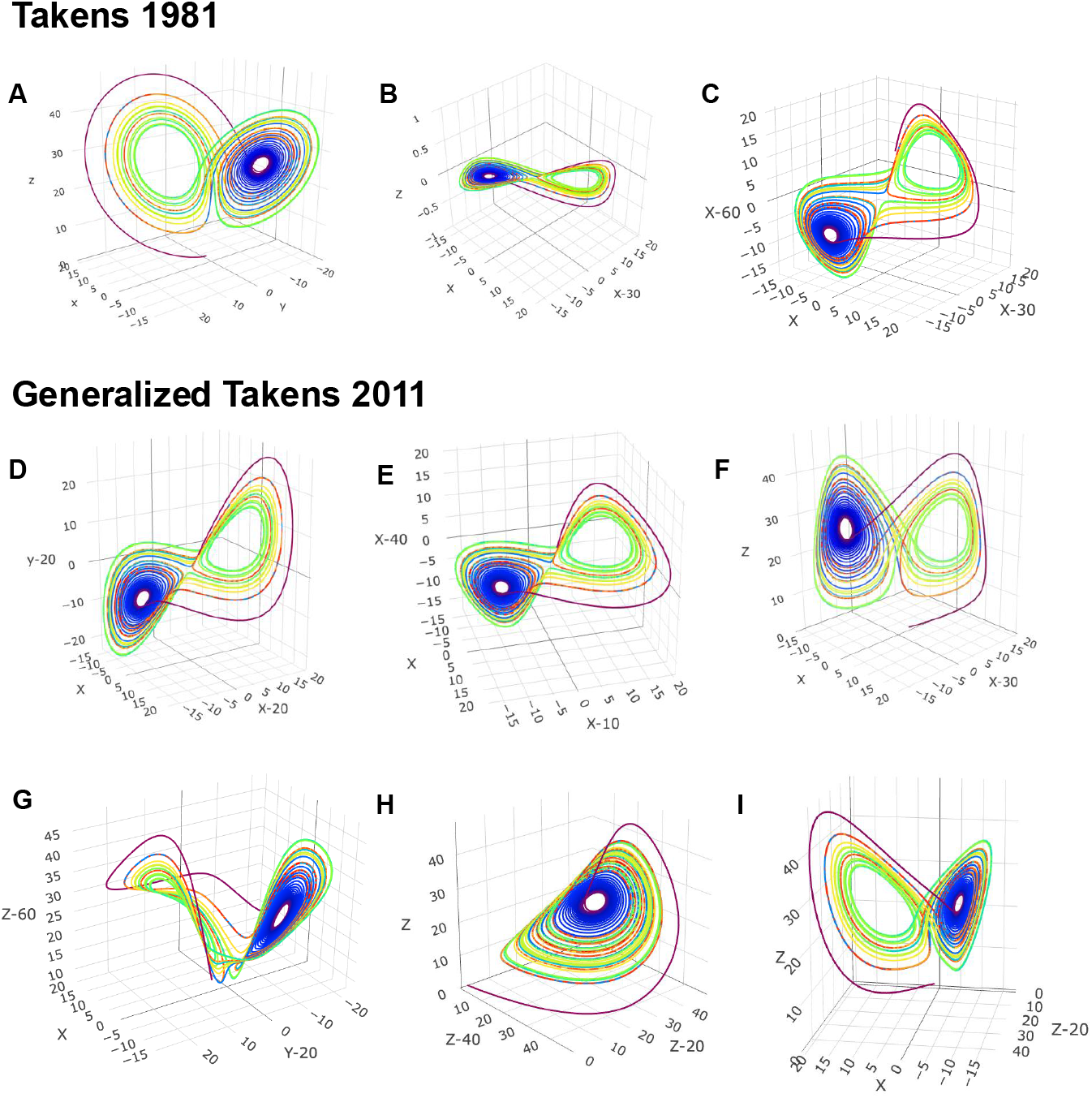
Takens theorem and generalized Takens embeddings. Takens’ 1981 theorem [28] showed that under generic conditions (measurements are coupled to all relevant degrees of freedom in the system) the attractor of a dynamical system can be reconstructed from a single observed time series, up to diffeomorphism. Panel A shows the native three-dimensional Lorenz63 attractor. If only the *X* time series is observed, a two-dimensional delay embedding using *X*(*t*) and *X*(*t* − 30) produces the projection shown in Panel B. Because this embedding lies in a plane, trajectories intersect near the two lobes, creating singularities and ambiguous states. Adding a second delay, *X*(*t* − 60), yields the three-dimensional reconstruction in Panel C, which resolves these crossings. The similarity between Panels A and C illustrates the diffeomorphic relationship: one manifold can be smoothly deformed into the other without tearing or folding. Deyle and Sugihara’s 2011 generalized Takens theorem [22], extended this result to mixed embeddings, in which delayed coordinates can be combined with contemporaneous observations of one or more variables. Panels D and F show examples using mixed coordinates composed of observed variables and time-delayed coordinates. Panel E illustrates that unequal delays, corresponding to different time scales, can also be used. Panel G further shows that embeddings may combine different variables with different delays. Mixed embeddings also help overcome known failure modes of the original Takens construction. In the Lorenz63 system, the left and right lobes of the butterfly attractor are symmetric about a fixed point on the *Z* axis. As a result, a delay embedding based only on *Z* collapses the two lobes into a single attractor and fails to preserve diffeomorphism Panel H [58]. Replacing one of the *Z* delays with *X*, or with a delay of *X*, restores the correct topology and resolves this degeneracy Panel I.

**Fig. E4:**
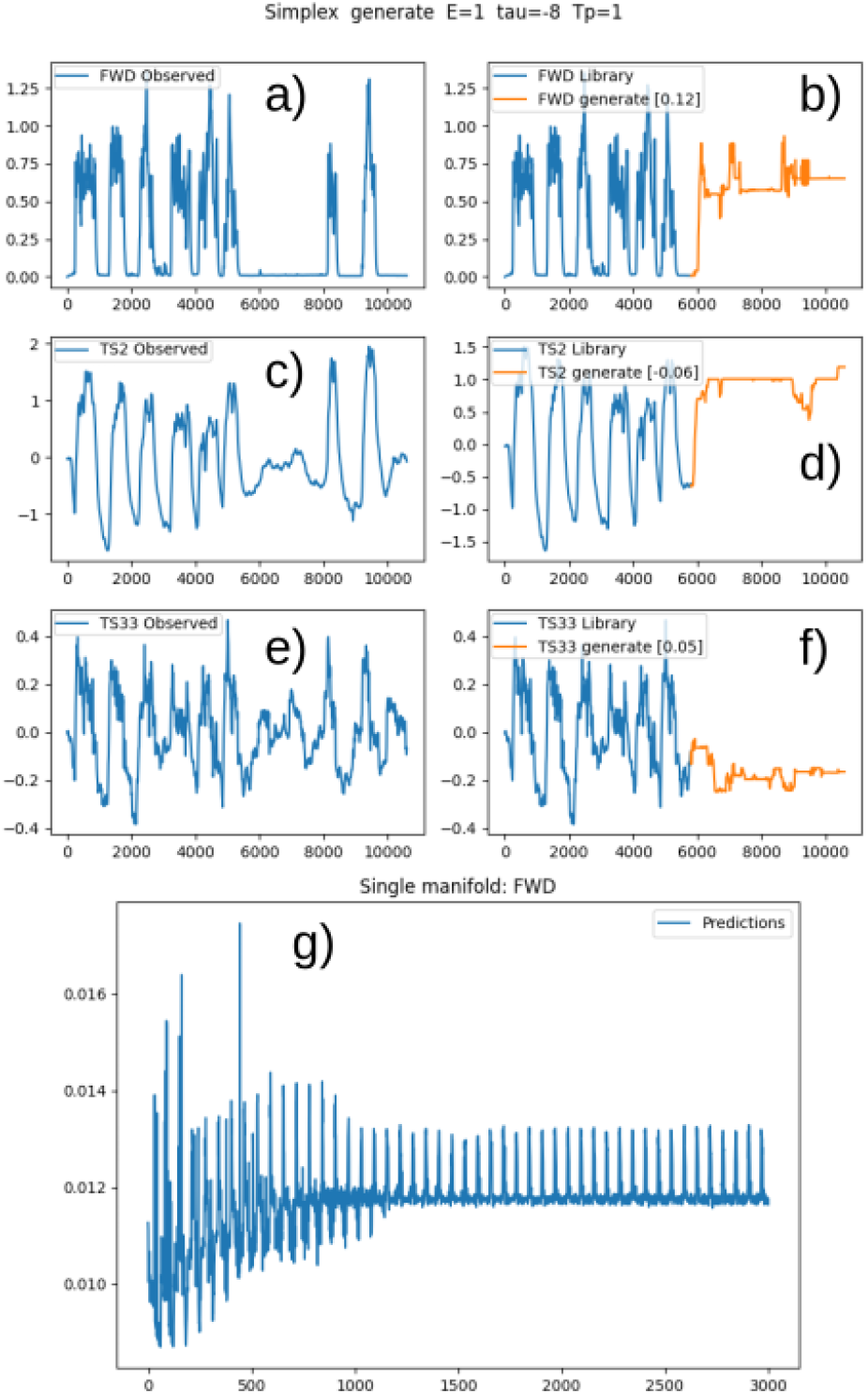
Generated time series of fly FWD motion without multiscale information (E=1, right column), and without network structure (single manifold, bottom panel). a) Fly forward motion (FWD). b) Forward motion generated dynamics without multiscale information starting at index 6000. c) Fly TS2 region neural activation. d) TS2 generated dynamics without multiscale information starting at index 6000. e) Fly TS33 region neural activation. f) TS33 generated dynamics starting at index 6000. g) Fly forward generated time series from a single manifold (no network).

## Supplement F *Drosophila* generated time series

Here we demonstrate good fidelity of GMN generated neural activity of arbitrary nodes in the network to observed time series substantiating holistic, multivariate time series simulation of GMN.

**Fig. F5:**
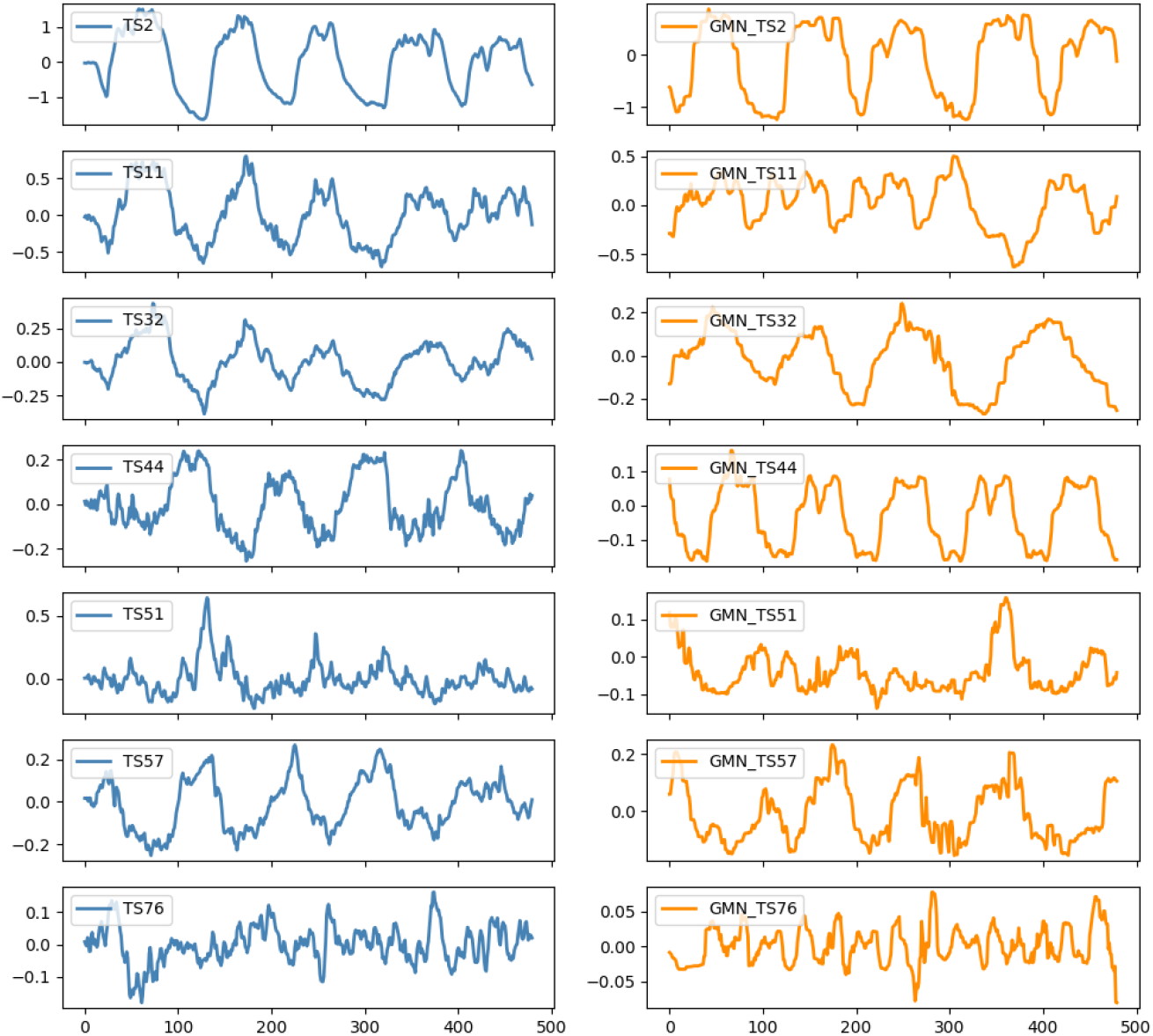
Left column: observed neural activations of *Drosophila*. Right column: GMN generated time series.

## Supplement G Reservoir computers

Reservoir computing (RC) is a vibrant field with recent reviews provided in references [38, 59]. Recognition of the echo state property [60] gave rise to echo state networks (ESN) which were celebrated for their ability to accurately generate short term chaotic dynamics [61, 62] as well as generating long term dynamics consistent with the underlying manifold. The reservoir represents a high–dimensional random embedding and as such there is an affinity to Koopman operators. A central tenant of Koopmanism is that within an infinite dimensional space a linear approximation to any nonlinear dynamic may be found, and RC have been shown to be Koopman operator approximators [63].

However, as demonstrated in reference [61] the output is highly sensitive to the reservoir structure and parameters. Thus one trades ease of construction without reservoir training with optimization of reservoir structure and parameters. Nonetheless, the ease with which an ESN can be configured and trained (conventionally, only the output layer is trained) coupled with recognition that they are universal approximators [33] has fueled their application.

Recent developments focus on expanding the capability and information content of the trainable output layer to better represent nonlinear dynamics [39, 64] or reduce reservoir dimensionality [65]. While these approaches are demonstrated to improve RC performance under specific conditions, conceptually one can question why a universal approximator needs nonlinear output modulation or storage of dynamical information in the output layer. This is likely to introduce fragility and it is perhaps better to represent dynamics in the reservoir. Indeed, it is recognized that RC require substantial warmup (training) to capture attractor dynamics, and even though next generation reservoir computers (NGRC) address this by adding output nonlinearites, they are critically sensitive to the choice of readout nonlinearity [39].

## Supplement H GMN Interaction and Node functions

Table H1 lists candidate interaction functions to discover the manifold network.

**Table H1:** GMN Network interaction functions.

| Purpose | Functions |
| --- | --- |
| Nonlinearity | $\rho_{\Delta}$ [21], $MI_{NL}$ [20], S-map [19] |
| Shared information | Simplex cross map [27], Mutual information (MI) |
| Shared dynamics | Convergent cross mapping (CCM) [18] |
| Separable dynamics | CCM with minimum MI |
| Separability | Clustering, PCA |
| Linear dependence | Correlation |

Table H2 lists candidate generator functions currently available in the `dimx`

**Table H2:** GMN Node generator functions.

| Function | Reference |
| --- | --- |
| Simplex | pyEDM Simplex |
| S-map | pyEDM SMap |
| k-nearest neighbors | sklearn neighbors.KNeighborsRegressor |
| Support Vector Regression | sklearn.svm SVR |
| Linear | sklearn.linear_model LinearRegression |

